# Goal directed closed-loop training of network bursts in cortical cultures produces improved performance and error correction over days

**DOI:** 10.64898/2026.10.15.759184

**Authors:** Wesley Clawson, Caitlin Grasso, Sydney Palmer, Serena Meng, Nathan Wu, Trevor Sullivan, Michael Levin

**Affiliations:** Allen Discovery Center at Tufts University, Medford, MA 02155, USA; Harvard University, Cambridge, MA 02138, USA; Google, Chicago, IL 60607

**Keywords:** diverse intelligence, learning, neuroscience, closed-loop control, embodiment

## Abstract

Dissociated neural cultures are living systems with no evolutionary history in their current form, which makes them a testbed for asking what goals neural tissue can learn to pursue once it is removed from its evolved body. It remains unclear which features of network activity such a culture can be trained to control, and whether that training is retained from one day to the next. We built an open-source closed-loop platform, the Holistic Agential Learning platform (HAL), that detects a network burst on a high-density microelectrode array, estimates its spatial origin in real time, and applies a minimal reinforcement rule. Here, a burst that originates off the goal side triggers global random stimulation that persists until the next burst. Over days of training, rat cortical cultures (n=10) increased the fraction of bursts that initiated on the goal side. This gain accrued between sessions, with little change within a session, a consolidation-like pattern that we interpret as a statement about timescale while the mechanism remains untested. The burst origin moved toward the goal side, while the burst peak and the repertoire of propagation directions were preserved. Trained cultures also recovered from incorrect states more reliably over days, while the rate at which they entered those states was unchanged, and this recovery effect was the most robust difference from unstimulated controls (n=6). Cultures that began on either side of chance converged to a similar level of performance. Damaged cultures (n=6) with comparable burst statistics showed neither the across-day gain nor the consolidation-like pattern. Training reweighted the activity patterns each culture already had, and we saw no new ones, which we read as a simple form of niche construction in which the tissue keeps a trace of its history that biases what it does next. These results identify burst initiation and error recovery as trainable variables in a novel living system, and they support training the tissue rather than engineering the machine.

## Introduction

Biological systems have a remarkable ability to adapt, or learn, relationships between themselves and environmental cues. How these systems do this is an open question, but better understanding the many forms of learning is of extreme relevance to applications in bioengineering and biomedicine [1, 2]. The trouble lies within the complexity of biology, typically with evolved systems building upon evolved systems, so much so that ‘building’ learning from the ground up seems an impossible task. It does seem though that the many scales found in biological systems understand how to work together, including learning new behaviors without disrupting others. Therefore, methods that allow the system to set goal states without restricting how those goals are met are experimentally attractive. Operant conditioning – learning a desired behavior through a system of rewards and punishment – is one of those methods [3]. While typically done to train behavior in animals, it has seen less use in synthetic biology as a way to train novel behaviors when systems are uncoupled from their normal sensory and processing loops by ‘embodying’ them in closed-loop experimental systems [4]. A dissociated culture on a microelectrode array is one such system. It is a living form with no evolutionary history in its current configuration, which makes it a useful testbed for asking what goal-directed competencies neural tissue retains when its evolved body and environment are removed [4].

While the field of neural computation is on the rise [5], it remains unclear what forms of learning (habituation, sensitization, goal-directed, etc.) are possible with the basal mechanisms of neural tissue once it is removed from the intact brain and its evolved anatomy, sensory streams, and motor pathways. This is a wide and largely open gap, however there is recent work on goal-directed learning in cortical organoids . Dissociated cortical cultures, grown in 2D sheets, are much more experimentally tractable while retaining key physiological properties of neural tissue in vivo.

Grown on a microelectrode array, tissues form large networks outside of predefined function -yet generate rich, structured, spontaneous activity, which allows for the investigation of how learning arises and is maintained from network dynamics themselves rather than from task-specific wiring [6–8]. A growing body of work suggests that learning-like adaptation does emerge in such simplified systems [9, 10], and interestingly that the capacity for trained, retained change may not be exclusive to brains at all [4, 11]. What remains unsettled is not whether cultures can change with experience, but which network variables can be trained, i.e. controlled by the tissue through feedback, and whether that change can be held ‘in tissue’ from one day to the next as a type of memory.

The foundational work of Shahaf and Marom demonstrated that a random network of cortical neurons could be trained to produce a chosen response: stimulation was applied until a desired activity pattern appeared on a target electrode, at which point it was withdrawn, so that stimulus cessation served as feedback [12]. Over repeated cycles the network needed fewer stimuli to produce the target, yielding clean learning curves, and the authors framed the process as a search through the network’s solution space of activity patterns, stabilized by feedback and requiring no explicit reward, supervision, or predefined representation. Modeling later hypothesized the underlying principle, learning by stimulation avoidance, which treats stimulus removal as a control signal and reproduces the stabilization behavior of the original experiments from simple plasticity rules combined with feedback [13]. Empirically, adaptive, feedback-driven protocols were shown to enhance plasticity and learning in culture relative to fixed-protocol stimulation [14, 15]. Steve Potter’s lab performed the embodied versions of the same idea, from which we took great inspiration, where a culture’s activity controls a simulated body and feedback returns through its inputs, which extended the demonstration from abstract responses to goal-directed behavior, and now its shown more recently in networks that learned to play a simple game within minutes [16–18].

Whatever the readout, the activity these networks produce is not only temporally but spatially structured, and is easily examinable in 2D. Network bursts do not ignite uniformly across the arrays they are grown on, but emerge from stable initiation zones and propagate as stereotyped, repeatable waves and motifs so that each culture expresses a limited repertoire of spatiotemporal patterns that have been described as attractors and even used as a functional fingerprint of the network [7, 19–25]. Stimulation can bias burst rate and action potential propagation [26, 27], but the initiation zones themselves have generally been treated as intrinsic, fixed by morphology, excitability gradients, or noise rather than by experience.

This leaves a two-pronged gap. First, as Shahaf and Marom proposed, learning in culture can be framed as a search through a solution space. That space has so far been defined temporally by asking what response follows what stimulus, and when. Whether training can also act on the spatial organization of activity, where bursts originate rather than only when they occur, is untested. Put directly: can closed-loop reinforcement move a burst’s initiation zone toward a goal? Second, learning has almost always been read out as a change in the level of a response, a rate or probability of a correct output measured at a point in time. Rarely has it been read out as a change in the dynamics by which a network moves between correct and incorrect states. For example, if a culture is punished for landing in a given state, or demonstrating incorrect behavior, two things could happen, and they are not mutually exclusive. The network could learn to avoid the actions that lead into that state, or it could learn to leave the state more reliably once it arrives there. The first is a change in what the network does and the second is a change in how it recovers. Distinguishing them, and testing whether training reshapes recovery time itself rather than just the odds of needing to recover, is a question about ongoing state-transition dynamics, not a fixed stimulus-response association [28, 29].

One important feature of learning in a system is that training, such as in operant conditioning or goal directed learning, produces a reliable change in behavior on a time scale longer than the training itself. Critically, the closed-loop and embodied demonstrations to date, including networks that learn to play a simple game within minutes [17], have shown acquisition only within a single session. To our knowledge no work on tissue culture has shown a task held with sustained competency from one day to the next, nor a consolidation-like effect in which performance grows more across days than within a session. Whether a cultured network can hold and build on a learned objective over days, rather than only acquire it in the moment, is a question of high importance given the rising interest in bio-hybrid computation and intelligence [30]. Additionally, if network states can be trained to be altered on long time scales, the applications to biomedicine are immense as we could experimentally train and understand long-term sensitization or breaking of habituation to drugs, or training tissues to alter diseased states.

Here we address these knowledge and capability gaps. We designed an open-source, closed-loop platform (named the Holistic Agential Learning platform, HAL) that detects each network burst on a high-density CMOS microelectrode array (HD-MEA) [31], estimates the spatial origin of the burst in real time, and delivers a minimal reinforcement rule: a burst that originates on a goal side produces no stimulation, while a burst on the wrong side triggers a global electrical stimulation at random intervals that persists until the next burst. Using this paradigm, we find that trained cultures show a shift of the origin of their bursts toward the goal side across training, moving from early to late training sessions, while unstimulated controls show no such directional shift. This spatial reorganization is a movement of the initiation site rather than of the burst peak, and it leaves the repertoire of propagation directions that existed before training intact. Alongside this spatial signature we find not only do cultures burst more preferentially from the goal side, but that trained cultures become better at recovering from incorrect states, without a matched change in how often the network slips into them. Overall, error correction improves while the rate of error itself does not. The gains in both the goal and recovery accrue across days more than within sessions, in a consolidation-like manner, and these increases depend on each culture’s intrinsic spatial bias before training. Finally, we attempted to train damaged cultures that appear electrophysiologically like healthy cultures and found no trace of learning or error correction. All together, we extend the application of closed loop, goal directed learning by examining two genuinely distinct dimensions the culture can organize to control -a spatial one, where activity emerges, and a dynamical one, how reliably the network recovers after an error.

These results carry two conceptual additions to current tissue learning and biohybrid computing literature. First, burst initiation zones, long treated as fixed features of a culture’s wiring, are plastic and learnable, which bridges the temporal learning literature with the spatial dynamics literature and positions even minimal, unstructured networks as systems capable of multi-dimensional adaptive control. Second, goal directed learning can in fact build consolidation-like memory in tissue culture without the need of complex mathematical decoders and encoders in the closed-loop process [5], although they can assuredly be useful in such designs. To this second point, we believe this protocol works as a proof of concept for a particular paradigm of tissue-computer interface design in which we drive learning into the tissue itself, rather than sophisticated input-output machines. Our reinforcement rule is a simple, global signal whose only structure is based on burst origin. Therefore, we view the reorganization we observe as produced by the tissue alone and not a reorganization of a complex machine learning system. While the machine learning techniques are wonderful tools, we propose this as an argument for training the tissue rather than engineering the machine, an approach that is critical for deeper understanding of closed-loop neuroengineering and for embodied or biohybrid systems built on living neural networks. We also offer a way to read these results together, which we return to in the Discussion. If recurring burst patterns are treated as agents and the slow variables of the substrate as their environment, then training that reweights existing patterns across days, with no new patterns added, resembles a simple form of niche construction in which the tissue’s own history biases what it does next.

## Results

### A closed-loop platform for reinforcing the spatial origin of network bursts

To ask whether learning in a dissociated network can act on the spatial dimension of tissue bursting activity, we built a closed-loop platform that reinforces individual network bursts in real time (Figure 1). E18 rat cortical neurons were dissociated and plated on a high-density CMOS microelectrode array (HD-MEA), and the HAL software suite was used to detect each network burst, estimate where it originated, and deliver feedback until the next burst arrived. Bursts were detected in real time using a sliding window of 200 frames (∼20 ms) and characterized by percentage of active electrodes of the recorded set. Further details can be found in the methods section, but briefly, bursts were characterized by (1) onset: >15% active electrodes, (2) origin: the average x position at onset, (3) peak: average x position at max % active electrodes in the burst, and (4) offset: when % active electrodes fell below the burst threshold for 50 ms. Each burst was classified as Left or Right according to whether the detected origin electrodes fell to the left or right of the average x coordinate at burst peak (Figure 1C, D). Our reasoning was that if the tissue grows homogenously across the MEA, the burst peak was physical midline of the MEA as burst typically span the entire tissue in the x direction. However, if a tissue grew only on the left 3/4ths of the MEA the burst peak would shift to the left. Our overall aim was that the tissue, not an input/output controller, was left to find a solution. Therefore, HAL defined a goal side for each trained culture and applied a deliberately minimal rule: a burst on the wrong side triggered an aversive, spatially distributed random stimulation that persisted until the next burst (punishment through its subsequent cessation) and a burst that originated on the goal side produced no stimulation (∼reward). In the presented data, we set the goal as left bursts for all trained cultures. Stimulating electrodes were chosen at random in the four corner quadrants of the array/tissue, with the intent that the aversive signal was global rather than a targeted, neuron-level instruction (Figure 1B). Each experimental day comprised a 20-minute pre-training baseline, 20 minutes of training, and 20 minutes of post-training recording. Due to software limitations at the time of study, we analyze activity from the unstimulated pre-and post-training phases only. This, however, allows us to capture how well the cultures were able to steer themselves to the goal state between sessions rather than observing immediate response to stimulation. Cultures were trained once they settled into a baseline (typically after DIV21), and not all cultures were trained at the same age. Therefore, the results sessions are aligned by training day, called ‘Adjusted Training Day’, indexed from each culture’s first training day. In total 10 cultures were trained in this protocol with 6 unstimulated controls plated alongside the first trained preparation (Table 1).

**Figure 1.**
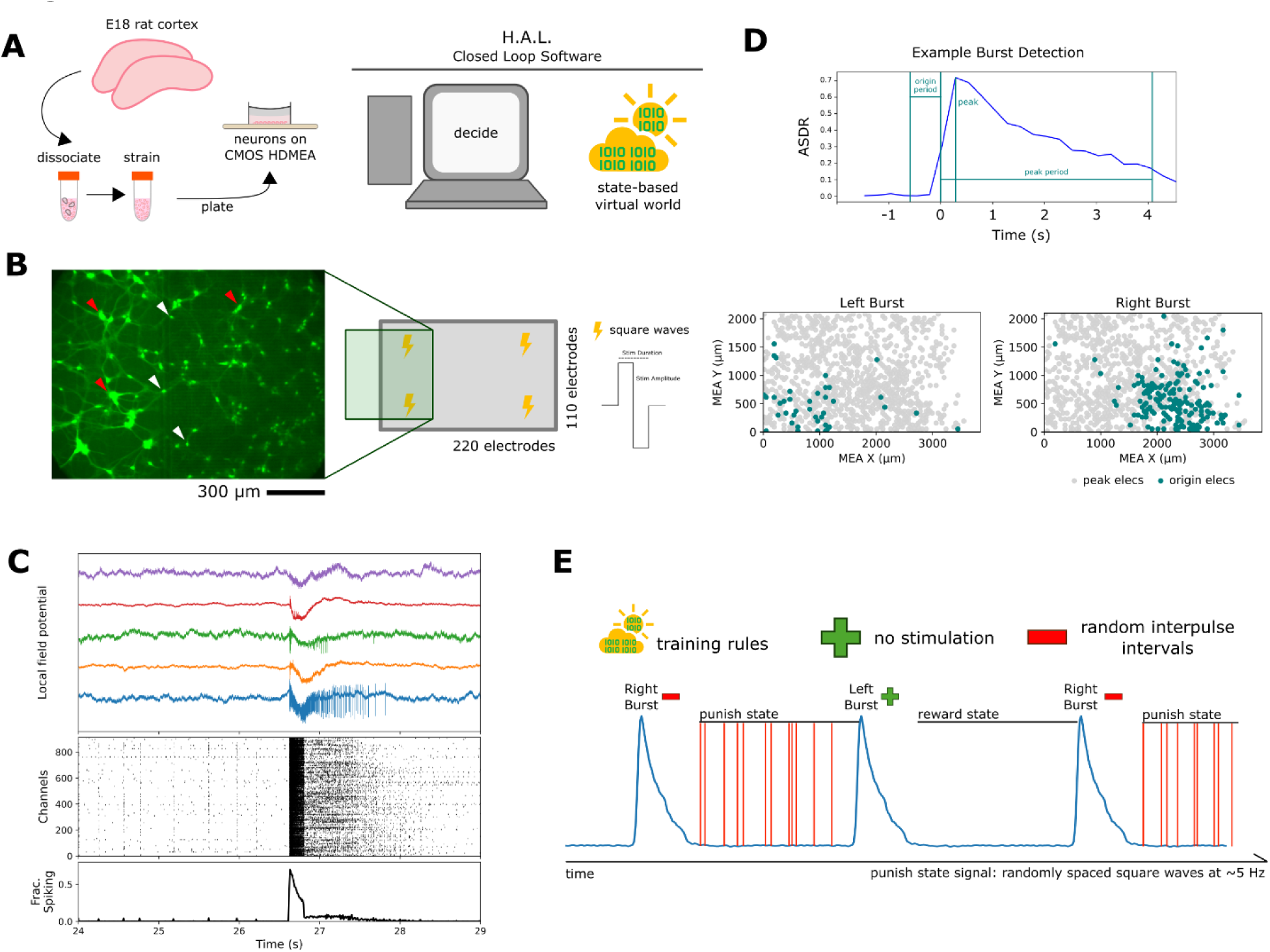
HAL Closed-loop reinforcement targets burst-origin dynamics at the tissue level. (A) Experimental workflow: E18 primary rat cortices are dissociated, passed through a cell strainer, and plated on a high-density CMOS microelectrode array (HD-MEA) connected to the HAL closed-loop system, which records network activity and delivers stimulation in real time according to predefined burst-detection parameters and world states (see Methods). Training rules are defined by the consequence of each burst: a correct burst yields no stimulation (reward), while an incorrect burst triggers an aversive random stimulation signal that persists until the next burst (negative reinforcement through its subsequent cessation). (B) Example neurons on a section of the array, stained with NeuO; red arrows mark small groups of neurons, white arrows mark individual neurons (scale bar 300 µm). Stimulating electrodes were chosen at random within the four corners of the array to deliver a global signal rather than to target specific neurons; stimuli were square pulses delivered simultaneously to all four electrodes. (C) Example traces of neuronal behavior (top), spiking activity across the dish (middle), and population firing rate (bottom). (D) Bursts are detected from the array-wide spike rate (ASDR); burst onset is defined as more than 15% of electrodes active, and HAL uses onset detection to set the next world state following burst offset. Each burst is classified as Left or Right according to whether its origin electrodes fall to the left or right of the burst peak. (E) Schematic of a training session promoting Left bursts: a burst originating on the right triggers aversive random stimulation until the next burst, a correct burst halts stimulation until the next burst, and a subsequent incorrect burst reinstates it. By convention the left side is the goal side, and all panels are oriented accordingly. Experiments comprise 20 minutes of pre-training baseline, 20 minutes of training, and 20 minutes of post-training recording.

**Table 1.** Culture cohorts. Cultures came from three preparations of E18 rat cortex. Controls were unstimulated sister cultures from preparation 1, recorded at an earlier age than the trained cultures. Damaged cultures were analyzed only for Figure 6. n is the number of cultures analyzed after the inclusion criteria; excluded were 2 stimRemoval cultures recorded only before the analyzed DIV range and 10 waveTraining cultures (7 with a nonstandard reward rule, 2 with a single recording day, and 1 with low activity).

| Preparation<br>(plating date) | Group | Dataset | Feedback rule | DIV<br>recorded | DIV<br>analyzed | n |
| --- | --- | --- | --- | --- | --- | --- |
| 1 (3 Jul 2024) | Trained | stimRemoval | Goal left; random<br>stimulation after right-origin bursts | 29 to 41 | 30 to 36 | 6 |
| 1 (3 Jul 2024) | Control | stimRemovalNull | No stimulation; coin-flip<br>goal side | 12 to 26 | 12 to 26 | 6 |
| 2 (3 Apr 2025) | Trained | waveTraining | Goal left; random stimulation after right-origin bursts | 32 to 43 | 34 to 42 | 4 |
| 3 (18 Nov 2025) | Damaged | burstTrainer | Goal left; random stimulation after right-origin bursts | 33 to 46 | 33 to 46 | 6 |

### Closed-loop reinforcement produces retained across-day gains consistent with consolidation

To test whether closed-loop training was associated with a change in burst origin, we tracked the fraction of bursts that originated on the goal side across training days (Figure 2A). We find that trained cultures increased their fraction correct within roughly two days and stabilized near 70-80% correct bursts. Comparing each trained culture on its first versus its last training day, performance increased significantly (Wilcoxon signed-rank test, W=5.0, p=0.019, n=10; Figure 2B). Controls, by contrast, did not improve over the same window (Wilcoxon W=9.0, p=0.844; Supplementary Figure S3). Controls have no real goal side as they do not undergo training and, by chance, their normal development may be read as a burst side preference. Yet, to measure fraction of correct bursts, a goal side must be chosen. We did not want to bias the control vs trained comparison in favor of training by choosing a goal side opposite to an innate preference, or bias against the training by choosing a goal side that aligned with preference. Therefore, we measure fraction of correct bursts in the n=6 controls as if the goal had been right or left, creating 2^6^ (64) possible cases (see Methods). In Figure 2A, the grey shading around the mean represents the variation across all possible outcomes for the controls, with the maximum and minimum being the ‘worst’ and ‘best’ cases. There is significant difference between average control and trained performance (Mann-Whitney U=51, p=0.022), but across the full set of 64 assignments the difference reaches significance in only 23% of cases, with a median two-sided p of 0.18. The per-culture effect size, the change in fraction correct from first to last day, was larger in trained cultures than in controls (Mann-Whitney U=51.0, p=0.022; n=10 trained, 6 control; Figure 2C). Therefore, we treat the effect size between trained and control as only partially significant, a best case rather than a robust difference.

**Figure 2.**
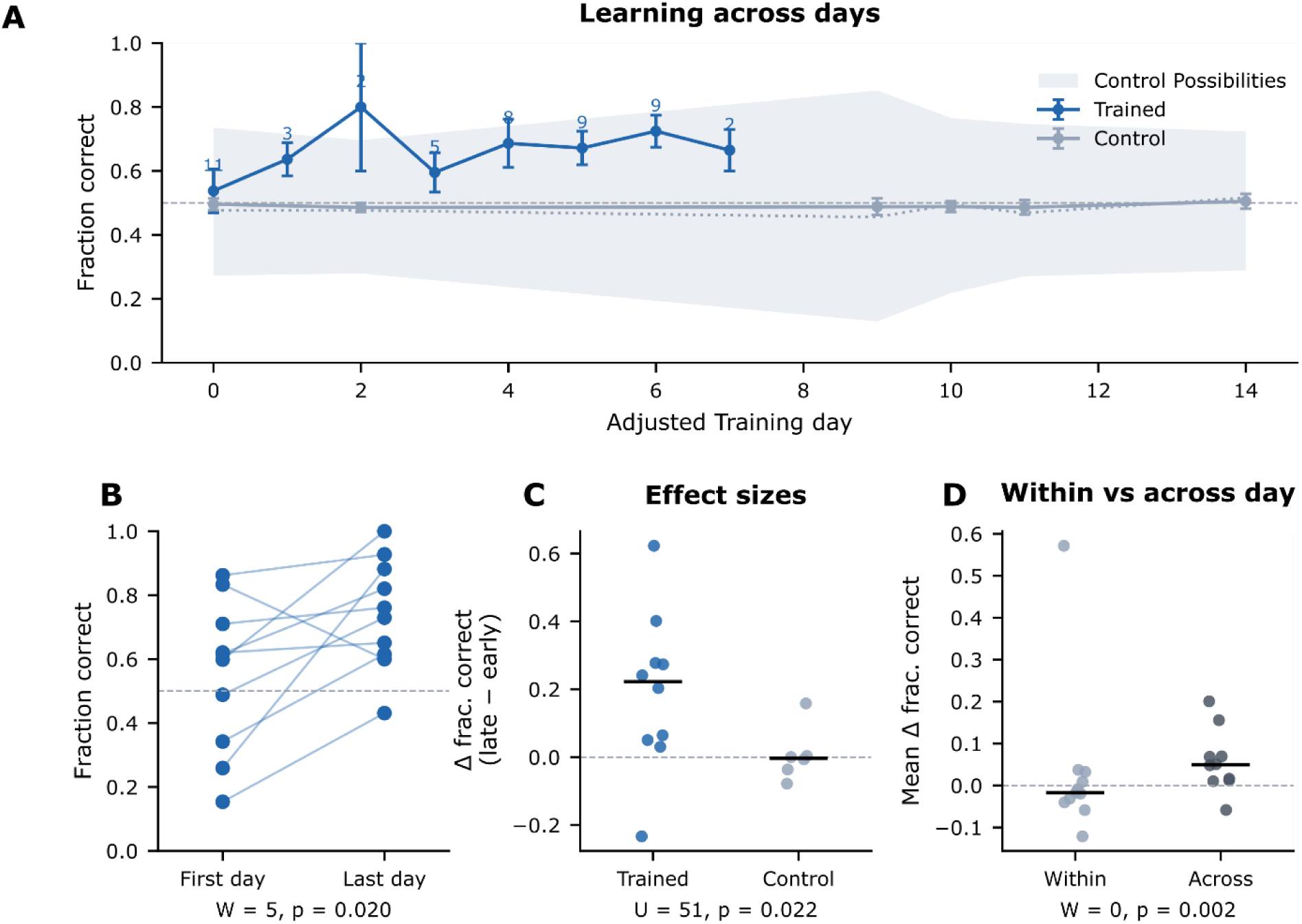
Closed-loop reinforcement drives across-day learning consistent with consolidation. (A) Fraction of correct bursts during the pre-training phase for trained (blue) and control (grey) cultures, plotted across adjusted training day, since cultures were not all trained on the same DIV. Trained cultures increase their fraction correct within roughly two days and stabilize thereafter; dots are the per-day mean and bars are SEM. Controls have no real goal side, so their fraction correct is averaged over 100 random coin-flip seed assignments to avoid accidental bias (see Methods); the shaded grey band shows the minimum to maximum range of control fraction correct across the 64 possible goal-side assignments. (B) Fraction correct for all trained cultures on the first versus last training day; performance increased significantly across training (Wilcoxon signed-rank test, W=5.0, p=0.019, n=10). (C) Effect size, the change in fraction correct from first to last day, trained versus control; trained cultures gained roughly 20% and more than controls on average (nominal Mann-Whitney U=51.0, p=0.022), but this contrast is not robust to the control coin-flip variance, reaching significance in only 23% of the 64 possible goal-side assignments (exact-null median p=0.18); we treat it as a supporting comparison and rest the learning claim on the within-trained tests in B and D (n=10 trained, 6 control). (D) Within-day versus across-day change in fraction correct for trained cultures; the change accrued between days far exceeds the change within a session (Wilcoxon signed-rank test, W=0.0, p=0.002, n=10), consistent with a retained between-session update; this analysis does not identify the cellular mechanism of consolidation.

We next asked when this change accrues, as traditional learning approaches search for learning across days/training. For each trained culture we separated the change measured within a single session (pre to post on the same day) from the change measured across days (post of one day to pre of the next) and found that the across-day term was significant (Wilcoxon W=0.0, p=0.002, n=10; Figure 2D). The across-day comparison between trained and control cultures was in the same direction but did not reach significance, consistent with an underpowered dataset (Mann-Whitney U=48.0, p=0.056; n=10 trained, 6 control; Supplementary Figure S3). Based on the within-trained tests, which do not depend on the control assignment, the paired first-to-last-day increase (Figure 2B), and the across-session increase in percent correct (Figure 2D), we interpret these results as directional improvement beyond natural developmental drift. We conclude that closed-loop reinforcement biases where bursts originate toward the goal side, relative to an unstimulated network. Additionally, learning under this paradigm is not a within-session short term change but a slow update that builds between sessions, a pattern we interpret as consolidation-like, but do not claim a demonstrated sleep or replay mechanism (see Discussion).

### Training improves how the network recovers from errors, not only whether it is correct

If the cultures had learned to direct their activity towards a goal state, we reasoned that the percent correct should increase, and that the way the network moves between reward and punishment states from burst to burst should also change. To test this, we treated the epochs between each burst as a state (state 0 on the goal side and correct, state 1 off the goal side and incorrect) and estimated the per-day transition probabilities between states in the pre-training phase, with each culture as the unit of analysis (Figure 3). Controls were again analyzed across the 64 possible goal-side assignments, so their transitions define a chance-recovery baseline. The fraction of time a culture spent in the correct state trended upward in trained cultures, but its first-to-last change did not separate from controls (Mann-Whitney U=43, p=0.18; n=10 trained, 6 control; Figure 3B). The recovery transition, the probability of moving from an incorrect to a correct burst, however, changed dramatically over training, with trained cultures becoming progressively better at leaving the incorrect state while controls stayed flat, and the first-to-last change in recovery was significantly larger in trained cultures (U=46, p=0.026 two-sided; p=0.013 one-sided for the a priori direction that training improves recovery; n=9 trained, 6 control; Figure 3C). This is the most robust between-group result in the study -it remains significant across 72% of the 64 possible control goal-side assignments (exact-null median two-sided p=0.034), unlike the effect-size comparison above.

**Figure 3.**
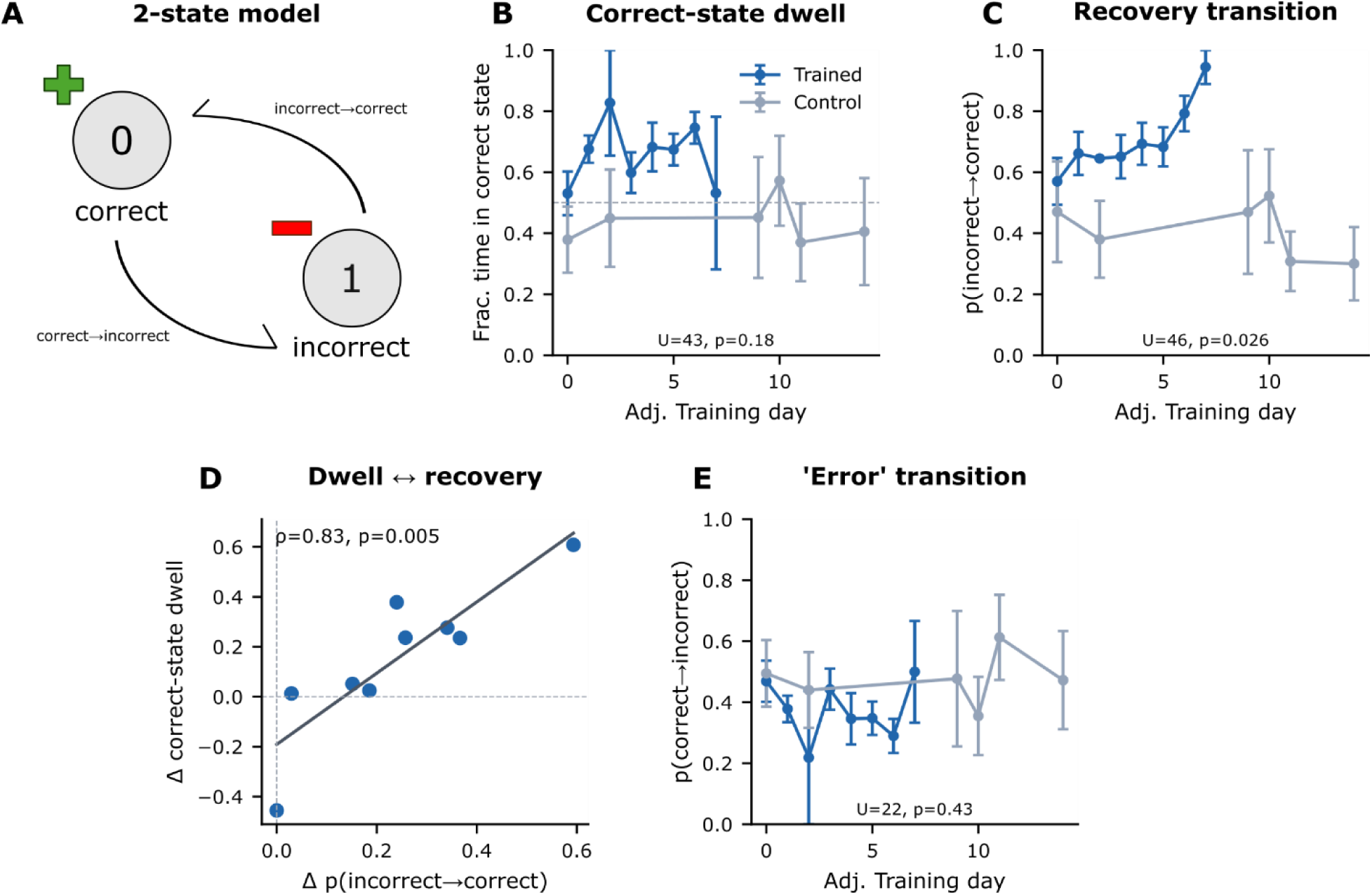
Closed-loop training preferentially increases recovery from incorrect burst states. Each burst is treated as a state (state 0, on the goal side and correct; state 1, off the goal side and incorrect), and we ask how training changes movement between states (pre-phase, unit of analysis is the culture; trained versus control). Control cultures are assigned the coin-flip counterfactual goal side, so their transition probabilities define a chance-recovery baseline. (A) Schematic of the two-state model and its transition probabilities. (B) Fraction of time spent in the correct state over training days; trained cultures trend upward, but the first-to-last change does not differ from controls (Mann-Whitney U=43, p=0.18; n=10 trained, 6 control). (C) Recovery transition, p (incorrect to correct), over training days: trained cultures increasingly climb back out of the incorrect state while controls stay flat, a significant difference in the first-to-last change (planned one-sided test for the a priori direction trained greater than control, U=46, p=0.013; robust to the coin-flip null, significant in 72% of the 64 possible control goal-side assignments, exact-null median two-sided p=0.034; n=9 trained, 6 control). (D) Across trained cultures, the first-to-last change in correct-state dwell and in recovery probability are strongly correlated (Spearman rho=0.83, p=0.005, n=9), with the regression line shown: cultures that become stickier in the correct state are the same ones that recover faster. Because dwell and recovery both encode being or becoming correct, this correlation is partly definitional, and we read panel D as internal coherence between the two measures rather than independent confirmation of the recovery effect. (E) Error transition, p(correct to incorrect), over training days does not differ between groups (U=22, p=0.43; exact-null median p=0.64, significant in 0% of assignments), so training improves recovery from errors without changing the rate at which the network falls into them, an asymmetry preserved across the coin-flip null. Recovery is reported as a single a priori directional test (panel C), with the error transition as its exploratory complement. A culture contributes a transition probability on a given day only if it had at least one transition from the relevant state; n=9 for the recovery test because one trained culture lacks a valid late-day estimate.

Conversely, the complementary error transition, the probability of moving from a correct to an incorrect burst, did not change with training (U=22, p=0.43; Figure 3E), so the effect is asymmetric -training improves recovery from errors without changing the rate at which the network falls into them. Across trained cultures, the change in correct-state dwell times and the change in recovery probability were strongly correlated (Spearman rho=0.83, p=0.005, n=9; Figure 3D), although because both quantities encode being or becoming correct, we read this as internal coherence between the two measures rather than as independent confirmation. We therefore identify a dynamical signature for this type of training -the trained network does not merely sit in the correct state more often, it recovers from errors more effectively over days.

### Training biases burst origins toward the goal side without shifting burst peaks

Having shown that training is associated with increased goal-side bursting and altered recovery dynamics, we next asked what spatial features of the burst changed. For each culture we computed the signed change, from early to late training, in the mean x-position of burst origins and of burst peaks, signed so that a positive value indicates movement toward the goal side (Figure 4D, see Methods). In trained cultures the origin moved significantly toward the goal side (one-sided Wilcoxon, W=46, p=0.032, n=10), whereas the peak did not move (W=18, p=0.375), and the magnitude of the origin shift exceeded that of the peak shift (paired Wilcoxon, W=3, p=0.010).

**Figure 4.**
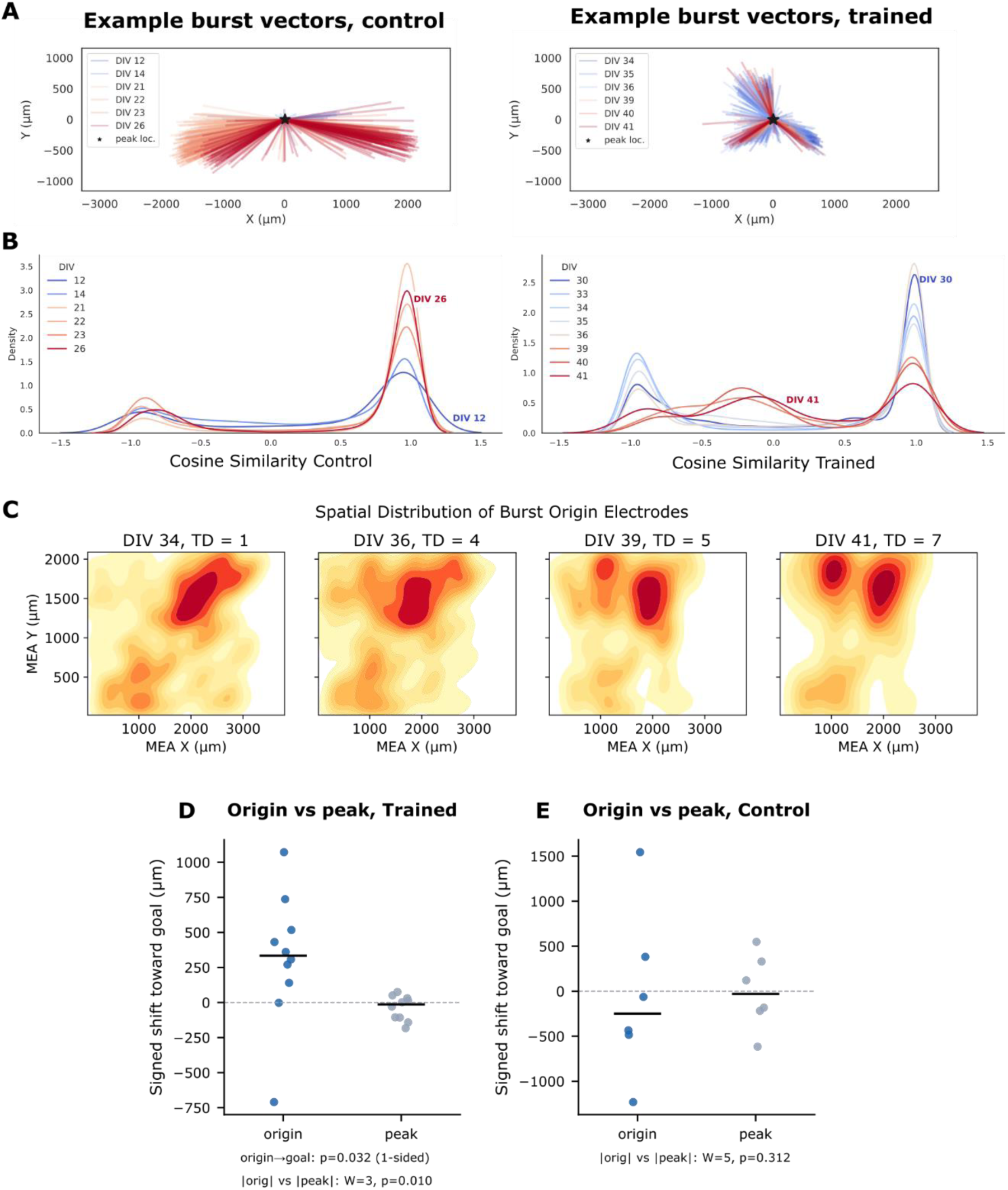
Training biases burst origins toward the goal side without shifting burst peaks. (A) Two example cultures, one control and one trained, tracked across DIV; each arrow points from a burst’s origin to its peak, with all peaks translated to the origin (see Methods). Both cultures develop structured, non-uniform origin distributions over development, illustrating that cultures naturally acquire burst-origin biases; in the trained culture these modes are redirected toward the goal side. (B) Density of cosine similarity across all burst vectors, pooled per DIV, for control and trained groups (minus one, opposite directions; zero, orthogonal; plus one, same direction). Both groups concentrate near plus one, and the diversity of propagation directions does not differ between groups (per-culture angular entropy U=34, p=0.80; mean cosine U=41, p=0.80; circular variance p=0.80): training reshapes where bursts originate without broadening or narrowing the repertoire of propagation directions. (C) Origin-electrode density for one example culture across training; warmer colors indicate higher density, and over training the origin mass shifts toward the goal (left) side. (D) Per-culture signed change in mean origin x-position and mean peak x-position from early to late training (each point one culture). Trained cultures move their burst origin toward the goal side (one-sided Wilcoxon, W=46, p=0.032) while the peak does not move (W=18, p=0.375), and the magnitude of the origin shift exceeds that of the peak shift (paired Wilcoxon, W=3, p=0.010, n=10). Control cultures show neither an origin shift (W=8, p=0.69) nor an origin-over-peak preference (W=5, p=0.31, n=6).

Control cultures showed neither an origin shift (W=8, p=0.69) nor any preference for moving the origin over the peak (W=5, p=0.31, n=6).

We then asked whether training reshapes the diversity of burst propagation directions or only their starting point. Comparing trained and control cultures per culture, we found no difference in the diversity of origin-to-peak directions by angular entropy (Mann-Whitney U=34, p=0.80), mean cosine similarity (U=41, p=0.80), or circular variance (p=0.80; Figure 4B, Supplementary Figure S5). The repertoire of directions is therefore preserved while its origin is displaced. We interpret this to mean that training shifts where activity initiates, not the set of trajectories the network is able to produce (see Discussion). We note that the between-group comparison of the raw origin shift, trained versus control, did not reach significance (Mann-Whitney U=41, p=0.26), so we rest this conclusion on the within-trained dissociation of origin from peak together with the control, rather than on the between-group contrast.

### Cultures converge to a similar performance despite disparate starting conditions

Not all trained cultures had the same shift in burst origin and we asked whether this heterogeneity was associated with each culture’s innate spatial bias. We classified cultures by whether the goal side aligned with their baseline-dominant burst direction (i.e. >50% of bursts before training day 1 were from the goal side; with-bias, n=6) or opposed it (against-bias, n=4), and followed fraction correct across training (Figure 5A, B). The two groups began far apart, with against-bias cultures starting well below chance, near 0.31, and with-bias cultures starting above it, near 0.71 (Mann-Whitney on first-day fraction correct, U=24, p=0.0095), yet they converged to a common band by the last day (around 0.79 with-bias and 0.67 against-bias; U=17, p=0.35). The larger raw improvement of the against-bias group may simply be its lower starting point rather than a signature of greater learning capacity. However, it is interesting that both groups on average converged to a similar % correct despite being a mix of tissues plated at different times with similar methods. We are careful here not to claim that training works better against a culture’s bias; but that cultures starting on either side of chance converge toward a similar level of performance (see Discussion).

**Figure 5.**
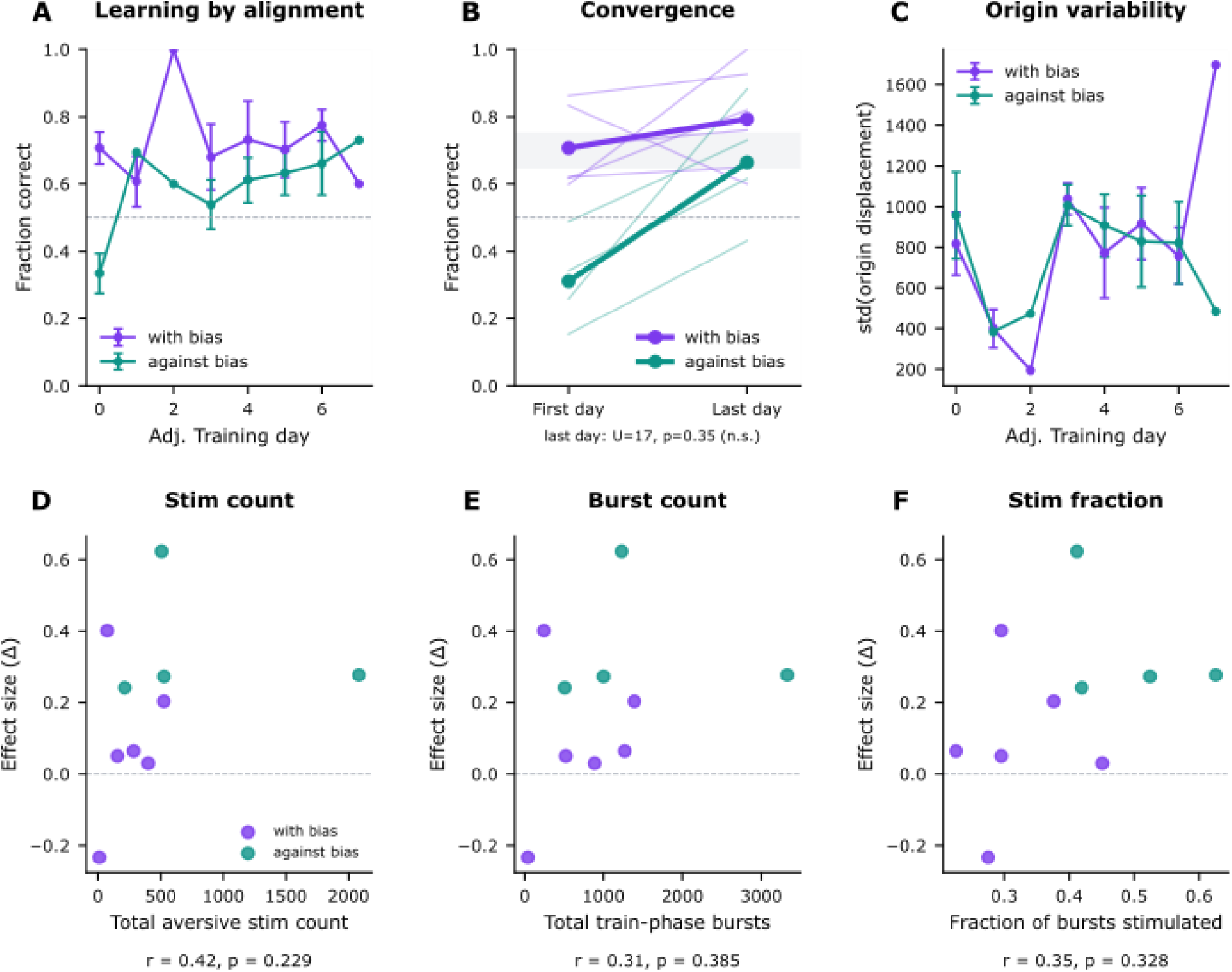
Training outcome depends on each culture’s intrinsic bias through its starting point, and is not explained by stimulation dose. Trained cultures are classified by whether the goal (left) side aligned with their pre-existing dominant burst direction (with-bias, purple) or opposed it (against-bias, teal); unclassified controls are grey. (A) Fraction correct over training days for with-bias versus against-bias cultures; against-bias cultures start well below chance and rise steeply, whereas with-bias cultures start above chance and change little. (B) First versus last day fraction correct by alignment group; the groups begin far apart (against-bias below chance, around 0.31; with-bias above it, around 0.71; Mann-Whitney U=24, p=0.0095) but converge to a common band by the last day (around 0.79 with-bias versus 0.67 against-bias; U=17, p=0.35). The larger raw improvement of the against-bias group therefore reflects its lower starting point rather than greater learning, and coin-flip controls reproduce the same with-versus-against gap. (C) Origin variability, the standard deviation of per-burst origin displacement per culture and day, over training by alignment group. (D to F) Per-culture effect size against three measures of stimulation or activity exposure: total aversive-stimulation count (D, Spearman rho=0.42, p=0.23), total train-phase burst count (E, rho=0.31, p=0.39), and the fraction of bursts that received stimulation (F, rho=0.35, p=0.33; n=10). None reaches significance, though these tests are underpowered and the three predictors are correlated, so we report no significant dose association rather than independence from dose.

A natural alternative is that the outcome simply reflects how much stimulation a culture received, as more errors means more stimulation, in line with claims from rate estimation theory [28, 29]. To address this, we related each culture’s effect size to three measures of exposure computed on the unfiltered train-phase data: the total aversive-stimulation count (i.e., number of biphasic pulses; Spearman rho=0.42, p=0.23), the total number of train-phase bursts (rho=0.31, p=0.39), and the fraction of bursts that received stimulation (rho=0.35, p=0.33; n=10). None showed a significant association (Figure 5D to F). These tests are underpowered and the three predictors are correlated, so we report no significant association with stimulation dose rather than independence from it; nonetheless, the learning we observe is not obviously a simple function of how much the network was stimulated. However, further studies need to be done to fully confirm this (see Discussion, Limitations).

### Healthy, but not damaged, cultures show within-group learning and consolidation

Finally, we asked whether spontaneous bursting is sufficient for this form of learning, or whether it requires intact network function, using a separate cohort of damaged cultures that continue to burst (Figure 6). We find that damaged tissues do not show a consistent increase in percent correct over training (W=8, p=0.69, n=6) as compared to the healthy tissue (Wilcoxon W=5, p=0.019, n=10), with their average percent correct decreasing from 0.59 to 0.56 (Figure 6C, D). We then examined if the consolidation-like effect existed in any form in the damaged tissue that may explain the lack of improvement. Again, we show that healthy cultures show no change in percent correct between pre-and post-training epochs on the same day, but have a notable increase between days (within versus across, Wilcoxon W=0, p=0.002). In contrast, the damaged tissues are extremely variable in their behavior, with a net decrease in percent correct within a single day, and no change across days (W=9, p=0.84; Figure 6E). The between-group statistics, by contrast, were underpowered and not significant (effect size healthy versus damaged, Mann-Whitney U=40, p=0.313; group-by-day interaction, cluster-robust regression p=0.111). However, the within group comparisons lead us to conclude that on average healthy cultures learn and demonstrate consolidation-like behavior, while damaged cultures do neither. Crucially, the damaged cultures were not silent, nor shockingly erratic. Their burst count, inter-burst interval, and burst duration were statistically indistinguishable from healthy cultures (burst count U=21, p=0.26; Figure 6F and Supplementary Figure S4). Spontaneous bursting is therefore necessary but not sufficient -the trainable reorganization we describe tracks intact network function across a collection of cells that is not explainable by generic electrical activity.

**Figure 6.**
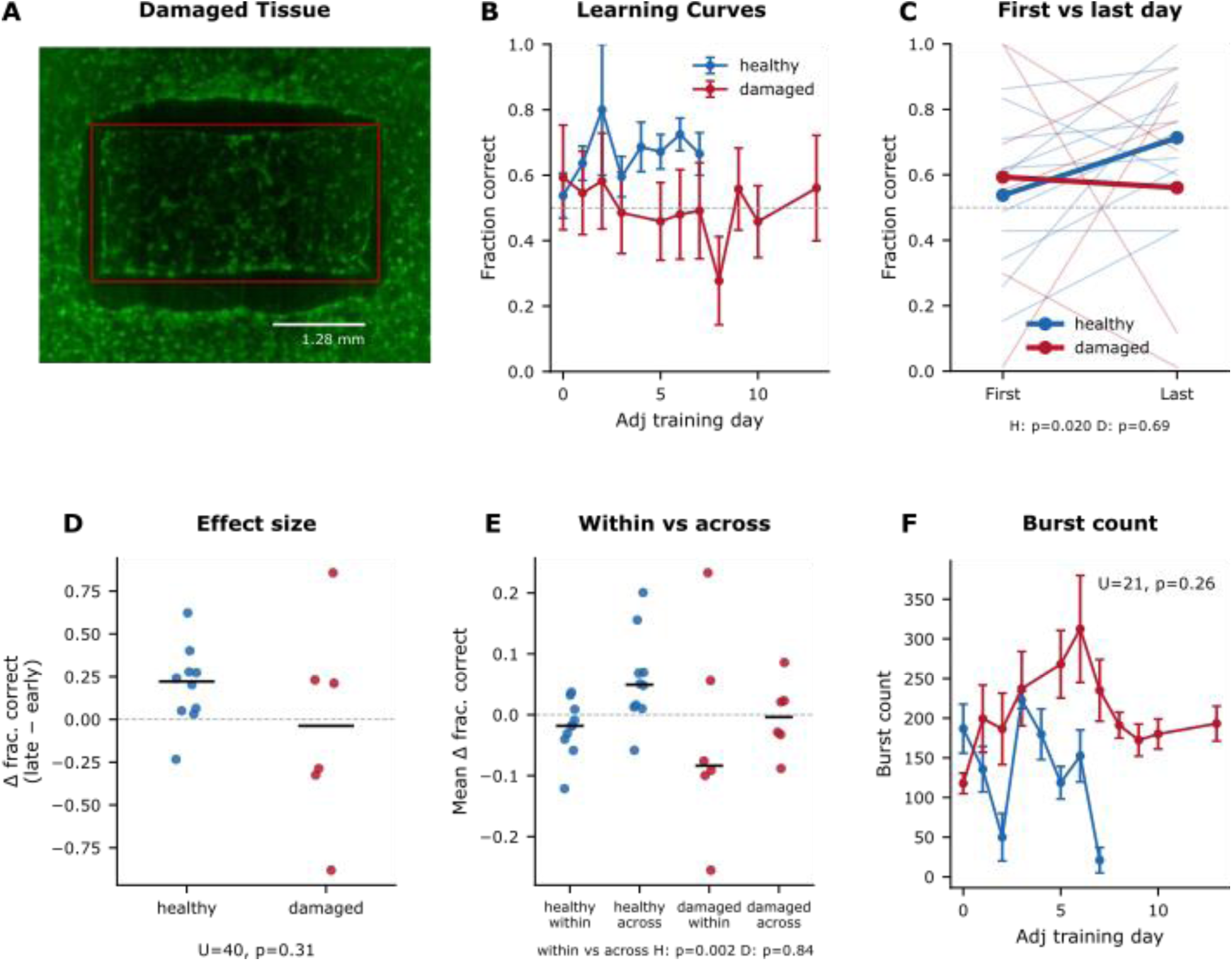
Healthy cultures show the retained training signature, whereas damaged bursting cultures do not. Healthy trained cultures (blue) are compared with damaged trained cultures (red); damaged cultures come from a separate dataset and are never pooled with the main cohort. (A) Representative microscopy of damaged tissue. (B) Fraction correct over adjusted training days: healthy cultures improve while damaged cultures remain near chance; a group-by-training-day interaction is not significant (cluster-robust regression, p=0.111) and the group means do not differ (Mann-Whitney U=44, p=0.301; n=11 healthy, 6 damaged; the healthy group in this panel includes one low-activity culture that is excluded from all other analyses), both comparisons underpowered, so the learning claim rests on the within-group tests in (C). (C) First versus last day fraction correct: healthy cultures improve significantly (Wilcoxon W=5, p=0.019) whereas damaged cultures do not (W=8, p=0.69; 0.59 to 0.56). (D) Per-culture effect size, healthy versus damaged (U=40, p=0.313; n=10 healthy, 6 damaged). (E) Within-day versus across-day change: healthy cultures show the consolidation signature (W=0, p=0.002) that damaged cultures lack (W=9, p=0.84). (F) Network burst count over training does not differ between groups (U=21, p=0.26); inter-burst interval and burst duration are likewise indistinguishable (Supplementary Figure S4). Damaged cultures thus retain broadly normal burst dynamics but lack the across-day learning shown by healthy cultures.

## Discussion

### Trained cell cultures exhibit goal-based learning and recovery from errors over days

We set out to ask whether a dissociated cortical network could be trained to reorganize its own activity through goal-directed, high-level feedback closed loop stimulation [32]. Critically, we wanted the tissue, treated as a whole, to be the controller for reorganization rather than targeted site or single neuron specific stimulations. We did this with the HAL closed-loop platform that rewards network bursts originating on a goal side (i.e. from the left of the array). We find that (1) trained cultures increase the fraction of bursts that initiate on the goal side over days, (2) that this gain is seen only across sessions rather than within them, (3) that it corresponds to a movement of the burst origin rather than the burst peak, and (4) that it is accompanied by an improved ability to recover from incorrect states. The effect varies with each culture’s intrinsic spatial bias and is observed in healthy but not damaged cultures; and in this dataset, it is not clearly explained by simple measures of stimulation exposure, such as in rate estimation theory. Together, these results extend the idea of learning in cultured networks along two axes – that cultures can be trained via goal-based learning approaches and burst site initiation is one parameter the tissue is capable of adapting towards a goal.

### Cultures learned not only to burst from the goal side, but to recover quicker from error

Early foundational work on learning in cultures demonstrated that dissociated cultures can learn and framed the process as a search through a solution space of activity patterns, stabilized by feedback when a desired temporal response appeared [8, 12]. That work, and the closed-loop and embodied studies that followed, measured learning almost entirely in the temporal domain, through response probability, latency, and rate [14–17]. In parallel, a largely separate literature established that network bursts are not spatially uniform: they emerge from stable initiation zones and propagate as stereotyped, repeatable motifs, such that each culture expresses a limited repertoire of spatiotemporal patterns sometimes described as attractors [19, 20, 22–24]. These initiation zones have generally been treated as intrinsic, set by morphology, excitability gradients, or noise. High-density recordings have since localized these zones to structural inhomogeneities in the network, with a few dominant initiation zones alternating as their local synaptic resources transiently deplete [21]. Our results attempt to join these two threads -we show that the origin point of bursts is itself a trainable variable, and that the network searches and stabilizes both the burst initiation site as well as how it responds in the presence of negative feedback (Figure 4). Importantly, this spatial reorganization is a displacement of the origin rather than an enrichment of the dynamic repertoire as the diversity of propagation directions is unchanged by training (Figure 4B, Supplementary Figure S5), so the network moves where its bursts begin without acquiring new trajectories. While it is indeed interesting that the burst origin is a trainable variable, these results hint that for dissociated tissue culture there may be limits on what can be trained through simple, closed loop experiments (no robust new dynamics). Two arms of rich future work would be to map what is adaptable and how best to map desired goal states to those trainable variables.

The state-transition result we believe adds an important piece to the picture (Figure 3). Rather than asking only whether a burst is correct, we asked how the network moves between correct and incorrect states, and found that training selectively improves recovery, the probability of returning to a correct burst after an incorrect one, while leaving the rate of errors unchanged. This asymmetry is informative: it suggests that what the network acquires is not a uniform suppression of incorrect activity but an improved capacity to correct itself, a property more naturally described in the language of rate estimation and ongoing adjustment than of a fixed stimulus-response association [28, 29]. We propose that spatial reorganization and improved error recovery may be coherent readouts of a common slow update, one indexing where the solution sits and the other indexing how reliably the network returns to it, while noting that the correlation between them is in part definitional rather than evidence for a single underlying process. Although the side-preference readout of Figure 2 is only partially robust, the improvement in error recovery is our most robust effect and strengthens across days, which is our central evidence that the network acquires something retained rather than merely tracking the reward in the moment. This in and of itself we believe is an important contribution to the field of in vitro learning, as many studies find that while cultures demonstrate a learned response during training, these effects cease outside of the training environment.

What exactly is retained, remains open. As in rate-estimation accounts of conditioning, a network that appears to learn a side preference may instead be tuning how it updates after errors, and separating these possibilities will require designs that vary the goal and the error structure independently, which we leave to future work.

### Across-day gains suggest a slow, consolidation-like learning process

A consistent feature of our data is that the trainable change is dominated by what happens between sessions rather than within them (Figure 2D; Figure 6E for the healthy cohort). The within-session component is small, and the bulk of the improvement appears as a difference between the end of one day and the unstimulated start of the next. We interpret this as a consolidation-like process -the network does not simply track the reward signal in the moment but updates slowly, in a way that is expressed after the stimulation is removed. We are cautious about the mechanism. Candidate processes include slow homeostatic or synaptic scaling [33, 34], structural plasticity accumulating between recordings [35–37], and stabilization during the long unstimulated intervals between sessions, but our data speak to the timescale of the change, not its biological substrate, and we do not claim a sleep or replay analogue. What the across-day dominance does establish is that the effect is a genuine, retained update rather than a transient response to ongoing stimulation, which is precisely why we characterized learning from the pre-training phase. Future studies in which HAL is coupled with automatic media sampling and testing, for example, could point to molecular pathways involved in this process.

### Reinforcement is linked to a culture’s innate spatial bias

Trained cultures did not all respond to training the same way, and the heterogeneity tracked each culture’s innate dynamical bias (Figure 5). Cultures whose baseline-dominant side already matched the goal began near ceiling and changed little, while cultures trained against their bias began well below chance and rose steeply, so that the two groups converged toward a common level of performance by the end of training. We are deliberately conservative in interpreting this convergence. Because alignment is defined by the relationship between the goal and a culture’s starting bias, the starting point is mechanically tied to the alignment label, and coin-flip controls reproduce the same with-versus-against gap; we therefore read the result as convergence from different starting points rather than as evidence that training is explicitly more effective against a culture’s bias.

We examined the stimulation dose to explicitly test the trivial explanation that learning is linked to simply receiving more stimulation. We show that a culture’s improvement is not significantly associated with the total amount of stimulation it received, the number of bursts it produced during training, or the fraction of bursts that were stimulated (Figure 5D to F). These analyses are correlational and underpowered, but we present them as ruling out an obvious confound rather than as a positive claim of dose independence, but they make it less likely that the learning we observe is a simple by-product of stimulation exposure. Future work will use yoked controls within the same dish, along with randomized total-stimulation-matched cultures, to better quantify these relationships, such as those shown in rate estimation theory [28].

### Damaged tissue does not demonstrate learning-like processes, despite similar activity

Dissociated neural cultures pose an interesting question – can neurons, torn from evolved cortical-typical circuits, retain some primitive functionality and express coordinated behavior necessary for learning. More simply, what kind of biological substrate can support dynamics, which then together perform simple goal-directed learning? We, fortunately, were able to assess part of this question by examining damaged cultures, although the small cohort limits direct between-group inference (Figure 6). They continued to burst, with burst counts, inter-burst intervals, and durations indistinguishable from healthy cultures, however they neither improved their fraction correct across days nor showed the across-day consolidation signature, while healthy cultures did both. We do acknowledge that this conclusion rests on within-group statistics rather than on a direct between-group difference, which our sample is underpowered to detect. These damaged cultures were not created on purpose, but by a fluke in the manufacturing process that caused the neurons to die along the edge of the MEA. We are therefore cautious to overclaim given the low n, but it is interesting that our embodied ‘tissue as the controller’ paradigm failed to produce meaningful learning signals despite appearing as an electrophysiologically healthy tissue. This raises a critical question of substrate – three substrates have similar high-level behavior, such as bursts, yet under the same goal directed learning only one shows significant change – what is the missing piece? It therefore remains a topic of future work to examine what aspect of the damaged substrate caused the loss of this function – was it a loss of neurons, a connectivity issue, or decoupled dynamics between the inner and outer sections of remaining tissue? Future studies could attempt to reproduce our damaged tissue, traumatic brain injury tissue culture models, or even other disease-in-a-dish models to investigate the necessary and sufficient consolidation-like mechanisms observed in the healthy tissue.

### Training as a simple form of niche construction

One way to read these results together is through stigmergy, coordination through traces left in a shared environment [38]. An action leaves a trace, the trace outlasts the action, and the trace biases what happens next. In a culture, recurring patterns of activity can be treated as the agents, and the slow variables of the substrate, such as synaptic weights, glial state, extracellular matrix, and the state of the closed-loop environment, as the niche that those patterns act on and inherit. Under this reading our data show a lower-order form of niche construction [39]. Training reweighted the patterns each culture already had, and we saw no new ones. This appears as a shift in burst origin over a preserved repertoire of propagation directions (Figure 4) and as a change in the transition probabilities between correct and incorrect states (Figure 3). The change also accrued between sessions (Figure 2D), which suggests that the trace is held in slow substrate variables that persist while the culture is unstimulated. A higher-order form, in which the culture’s own traces stabilize new patterns or remove old ones, would appear as a change in the repertoire itself, and we did not observe it. We did not measure any substrate variable directly, so we offer this reading as a framework for future experiments and make no claim about mechanism. The distinction it draws is measurable on high-density arrays: lower-order effects change transition probabilities over a fixed set of patterns, and higher-order effects change the set.

### Limitations

Our study had several limitations. Primarily that of a small n with three separate culture preparations yielding n=10 trained cultures, n=6 damaged cultures, and n=6 controls. We therefore relied heavily on within-group tests (pre-vs post-test), which were significantly different for measurements such as percent correct and error recovery, and noted when across group tests were underpowered. Second, the origin metric is a one-dimensional summary, the position of initiation along the x axis, and although it is sufficient to demonstrate that the origin moves while the peak does not, it does not capture the full two-dimensional structure of where bursts begin. Future work is planned to examine if cultures can train bursts towards quadrants, rather than sides, to see if cultures are capable of fine tuning the burst origins in both directions.

Our controls are unstimulated cultures, which does isolate the effect of goal-directed training from spontaneous drift but doesn’t separate the contribution of reward contingency from that of stimulation per se. We do see a slight correlation (rho=0.42) between total stim count and effect size of learning, but future work should include both yoked and randomly rewarded controls within the same 6-well dish. The yoked control would allow the decoupling of total stimulation and reward contingency, while the random stimulation would test if stimulation decoupled from the task had an effect.

Our cultures were plated as 3 separate preparations (Table 1, see Methods). Controls were plated alongside the first trained preparation but were recorded at an earlier age (DIV 12 to 26, versus DIV 30 to 42 for trained cultures), and the groups differ at baseline in inter-burst interval, burst size, and duration (Supplementary Figure S1); the same properties are shown across training days in Supplementary Figure S2. Burst properties change with culture age, and performance in closed-loop culture systems is known to depend on culture age and stimulation history [5]. While this is consistent with dissociated tissue prep [25], this does make comparing learning directly across the groups shaky, and the between-group effect size does not survive adjustment for baseline burst dynamics (Table T2). However, we find it interesting that despite these differences (1) the trained cultures all show a robust increase in percent correct and that this increase comes between days and (2) damaged cultures don’t demonstrate this at all, despite having activity not dissimilar to both trained and control.

### Outlook

The future of closed loop engineering around synthetic forms, such as dissociated primary tissue culture, is extremely bright. Here we show that cultures can self-organize their dynamics, and assumedly their substrate in the form of synapses, towards a spatial goal state and reduce error. The goal we set, a preferred side for burst initiation, is one that cortical tissue may have no evolutionary reason to pursue, and cultures that started on either side of chance still moved toward it. This is consistent with a classic operational definition of intelligence, the ability to reach the same goal by different means [40], although our convergence result carries the caveats noted above. Seeing it in a living form with no evolutionary history in this configuration supports treating goal-seeking as a property to be tested for in any substrate [4]. The design allows for self-organized behavior, and the 2D cultures are broadly more experimentally investigable through molecular and imaging techniques so future experiments can more reliably track what primitive functionality the neurons and networks are implementing to create change. These findings will be critical in bettering our understanding of the wealth of plasticity beyond synaptic, including how to control it. Tissue wide dynamics, such as bursts, are closely related to travelling waves in vivo, which are implicated in brain wide processes that support cognition [41–43]. Closed loop experimental design around travelling waves in cultures could either mechanistically understand these processes, or be used in neurocomputing processes, rather than trying to train them away or ignore them. A thrust of traditional cybernetics was a question of control, regardless of mechanism, and these closed loop systems may help us to explore how closed-loop systems in brain computer interfaces could heal, repair, or enhance cognition. Our results are some of the first that see long-term change, and we hypothesize that the cultured tissue is best treated as an agent to be persuaded rather than a machine to be programmed, engaged through reward and punishment in a manner closer to classical conditioning [44] than to direct control, and placed on an axis of agency and persuadability that spans biological substrates [4, 32]. Learning how to persuade a dish of neurons may prove a useful first step toward persuading an injured brain to repair itself.

## Acknowledgements

We acknowledge AFOSR grant FA9550-22-1-0465 and DARPA INSPIRE HR00112530152. This research was developed with funding from the Defense Advanced Research Projects Agency (DARPA), HR00112530152 INSPIRE. The views, opinions, and/or findings expressed are those of the author(s) and should not be interpreted as representing the official views or policies of the Department of Defense or the U.S. Government.

## Contributions

Study was designed and conceptualized by WC and ML. Initial HAL development was done by TS, NW, and WC with further refinement and development done by CG. Analysis was done by WC and CG. Tissue culture growth and training were done by WC, NW, SP. Writing was done by WC, SP, and ML. WC would like to acknowledge Maxwell Biosystems on advice on initial tissue plating work as well as insight on internal MxB software.

## Methods

### Cell culture and plating

Pre-conditioned microelectrode arrays (MEAs) were washed with sterile deionized water and coated with 0.07% Polyethyleneimine (PEI) (Sigma-Aldrich; Cat# P3143) in sterile borate buffer (ThermoFisher Scientific; Cat# 28341) for 1 hour at room temperature. The MEAs were washed with sterile deionized water and allowed to air dry for 1 hour before adding a secondary coating. Once dry, 0.02mg/ml murine Laminin (Sigma-Aldrich; Cat# L2020) in chilled sterile plating medium (NeuroCult (STEMCELL Technologies; Cat# 05711) supplemented with 0.5mM L-Glutamic Acid (Sigma-Aldrich; Cat# G1251), 12.5mM D+Glucose (Sigma-Aldrich; Cat# G8769), 0.5mM GlutaMax (Gibco; Cat# 35050061), 0.5% Penicillin/Streptomycin, and 2% SM1 Neuronal Supplement (STEMCELL Technologies; Cat# 05711)) was added to the MEAs and incubated for 1-3 hours in a CO2 incubator (37°C, 5% CO2). Primary and secondary coatings were plated with 50ul volume to cover the entirety of the array.

E18 Sprague Dawley Rat cortices were obtained from TransnetYX Tissue (TransnetYX, Cat# KTSDECX). The tissue was digested in pre-warmed 6mg/ml Papain dissolved in THRYVE Embryonic-CaCL (provided in TransnetYX, Cat #KTSDECX) for 10 minutes at 37°C. Cortices were returned to THRYVE medium and triturated with a fire polished salinized Pasteur pipette until completely disassociated (∼10 times). The cell slurry was then filtered through a 40um cell strainer, counted, and resuspended at 2.0 × 106 cells/ml in sterile plating medium. Laminin was removed from each MEA, leaving a thin film behind, and 50ul of cells (final 1.00 × 105 cells; ∼2,000 cells/mm2) were immediately plated directly on the array. Cells were maintained in a tissue culture incubator (37°C, 5% CO2). A 90% medium change with plating medium is performed on DIV 1. Beginning DIV 5, subsequent medium changes are 50% Cell Culture Maturation Medium (CMM) (BrainPhys supplemented with 0.5mM GlutaMax, 12.5mM D+Glucose, and 2% SM1 Supplement).

### Microelectrode array and recording

Closed-loop experiments used the MaxWell Biosystems MaxTwo system. The MaxTwo Well Plate contains 6 isolated arrays, each comprising 26,400 electrodes coated with platinum black on a 17.5 um pitched 3.85 x 2.10 mm^2^ rectangular sensing area. However, only 1024 channels can be recorded simultaneously at a maximum sampling rate of 10kHz per array. Recording electrodes were selected with the MaxWell Activity Scan assay, which surveys activity across the array by recording spiking in successive electrode configurations for 60 seconds each in order to scan the array for ‘Active Areas’. Spikes were detected as activity exceeding 5.5 times the root-mean-square (RMS) noise of the signal. Electrodes from the merged configurations were filtered by a firing-rate threshold between 0.1 and 0.2 Hz and an amplitude threshold of 20 µV, and the maximum number of electrodes was then drawn at random from this set to form the recording configuration. Four stimulation electrodes were selected from the recording set, placed in the four corner quadrants of the array (Figure 1) and chosen to be minimally involved in bursting so as to limit stimulation-induced activity.

### Closed-loop training with HAL

Training was run in real time through the Holistic Agential Learning (HAL) platform, which interfaces with the MaxWell Python and C++ APIs to configure the device and to process data and deliver stimulation online. On each iteration HAL read the incoming frame of spike information, appended it to a sliding window of 200 frames, and dropped the oldest frame. A burst was detected when activity in the window exceeded a threshold of 15% of the network; to be counted, activity had to remain above threshold for 100 ms and then fall below it for 50 ms, so that a brief dip during a single burst was not split into two. For each burst, HAL recorded the average x position of the active electrodes when the threshold was first crossed (the origin) and the average x position when activity in the window reached its maximum (the peak), and classified burst direction by comparing the origin x position to the peak x position.

Feedback depended on burst direction. A stimulation pulse was a biphasic waveform of 250 mV amplitude and 200 µs per phase. If the preceding burst was in the target direction, no stimulation was given (reward). If it was in the wrong direction, random stimulation was delivered as pulses at random intervals at a mean rate of 5 per second to the four stimulation electrodes (punishment), and continued until a new burst was detected. To avoid mistaking stimulation-evoked activity for a burst, the detector was silenced for 800 µs after each pulse. All cultures were trained for left bursting.

Trained cultures came from two preparations (n=6 and n=4 analyzed cultures; Table 1), and each culture was trained on upwards of 7 days depending on its development and viability. Each daily session consisted of a 20-minute pre-training recording (pre), 20 minutes of closed-loop training (train), and a 20-minute post-training recording (post). All learning analyses below use the unstimulated pre phase.

### Offline burst and epoch extraction

Recordings were processed offline into per-burst tables. Network bursts were detected from the array-wide activity using a 10 ms bin size, a 20 µV amplitude threshold, and a 15% network burst threshold, with a 50 ms window applied after stimulation to exclude stimulation-evoked frames.

For each burst we recorded its onset, peak, and offset times, duration, size as the percentage of active electrodes, and the x positions of its origin (electrodes active in the onset window) and its peak. The signed displacement was defined as dx = origin_x minus peak_x; bursts with dx less than zero were labeled left and bursts with dx greater than zero were labeled right, and bursts with dx equal to zero were dropped as ambiguous. Only network bursts of the standard type were analyzed. The inter-burst interval was defined as the time from the offset of one burst to the onset of the next, keeping only positive intervals.

For each trained culture the desired side was the side that avoided the aversive stimulation, and a burst was scored correct when its direction matched the desired side. The origin and the peak are distinct physical quantities, the origin being the centroid of the electrodes where a burst initiates and the peak the centroid of maximal array activity during the same burst, so a shift in one need not imply a shift in the other. Recordings were assigned to the pre, train, or post phase. Because cultures were trained at different ages, analyses were aligned by training day, indexed from zero at each culture’s first valid recording day. Learning analyses were restricted to each dataset’s valid DIV range (stimRemoval DIV 30 to 36, waveTraining DIV 34 to 42).

### Datasets, controls, and inclusion criteria

Trained cultures came from two datasets, stimRemoval and waveTraining. Control cultures were unstimulated sister cultures plated in the same preparation as the stimRemoval cultures and recorded at DIV 12 to 26 (stimRemovalNull; Table 1); because controls have no real goal side, a desired side was assigned as a deterministic coin-flip per culture, and for the learning curves the fraction correct was averaged over 100 random seed assignments to remove accidental bias. Damaged cultures came from a separate preparation (burstTrainer) and were analyzed only for Figure 6, never pooled with the main cohort. A small number of cultures that used a nonstandard reward rule were excluded. Cultures were included if they had at least 2 recording days and at least 10 train-phase bursts; this removed cultures that were never meaningfully trained. The sensitivity of the headline results to these filters is given in Table T1. Bursts that coincided with stimulation were excluded from analysis, except for the train-phase exposure measures in Figure 5D to F, which were computed on the unfiltered bursts so that stimulation counts were not undercounted. The unit of analysis was the culture throughout.

### Quantification and statistics

Fraction correct was the proportion of a culture’s bursts that originated on the desired side, computed per phase and day. Learning was quantified on the pre phase. For first-versus-last comparisons we took each culture’s first and last valid training day. Effect size was the change in fraction correct from the first to the last day. To separate within-session from across-session change (Figure 2D, Figure 6E), we compared the mean change within a day (pre to post) with the mean change across days (post of one day to pre of the next).

For the state-transition analysis (Figure 3) each burst was assigned a state (0 if correct, 1 if incorrect), and per culture and day we computed the dwell fraction (the fraction of inter-burst time spent in the correct state) and the transition probabilities between states, where p(incorrect to correct) is the recovery probability and p(correct to incorrect) is the error probability. A culture contributed a transition probability on a given day only if it had at least one transition from the relevant state. For the spatial decomposition (Figure 4) we computed, per culture, the signed change from early to late training in the mean origin x position and the mean peak x position, signed so that positive values indicate movement toward the goal side. Burst-direction diversity (Figure 4B, Supplementary Figure S5) was quantified per culture as the angular entropy, mean pairwise cosine similarity, and circular variance of the origin-to-peak vectors. For the bias analysis (Figure 5), cultures were classified as with-bias or against-bias according to whether their baseline-dominant side matched the goal side, and stimulation exposure was summarized as the total aversive-stimulation count, the total train-phase burst count, and the fraction of bursts that received stimulation.

Group comparisons used the Mann-Whitney U test for independent samples and the Wilcoxon signed-rank test for paired samples; associations used the Spearman rank correlation. Tests were two-sided unless an a priori directional hypothesis was specified, in which case the one-sided test is reported and labeled as such (the origin shift toward the goal in Figure 4D and the recovery improvement in Figure 3C). For the transition analysis, improvement in the recovery probability was a single a priori directional hypothesis and is reported as a one-sided test, with the complementary error transition reported as an exploratory two-sided comparison. The healthy-versus-damaged learning comparison (Figure 6B) additionally used a culture-clustered ordinary-least-squares model to test the group by training-day interaction. Because control cultures are assigned a coin-flip goal side, the trained-versus-control effect-size comparison (Figure 2C) was also assessed against the exact coin-flip null, enumerating all 64 possible goal-side assignments of the six control cultures and recomputing the test for each; we report the median p and the fraction of assignments reaching significance. Analyses were performed in Python (version 3.12) using NumPy, pandas, SciPy, and statsmodels.

## Supplementary Figures

**Figure S1.**
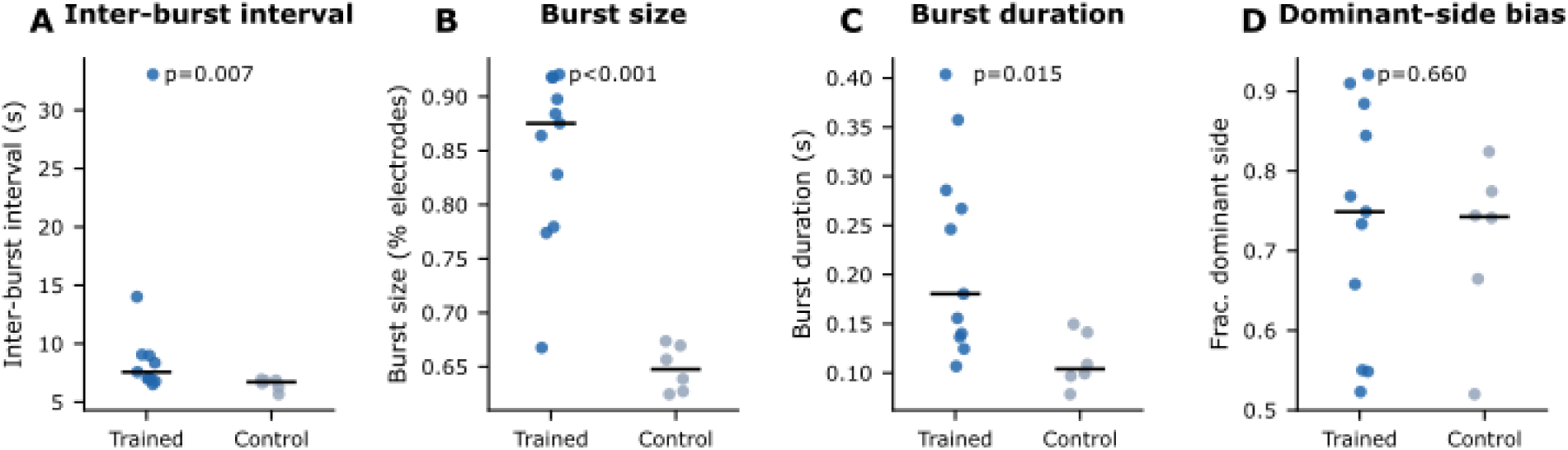
Baseline burst properties differ between trained and control cultures. Per-culture baseline measures: (A) inter-burst interval, (B) burst size as percent of active electrodes, (C) burst duration, and (D) fraction of bursts to the dominant side. Trained and control cultures differ at baseline in inter-burst interval (Mann-Whitney p=0.007), size (p<0.001), and duration (p=0.015), but not in dominant-side bias (p=0.660), indicating the groups are not perfectly matched at baseline.

**Figure S2.**
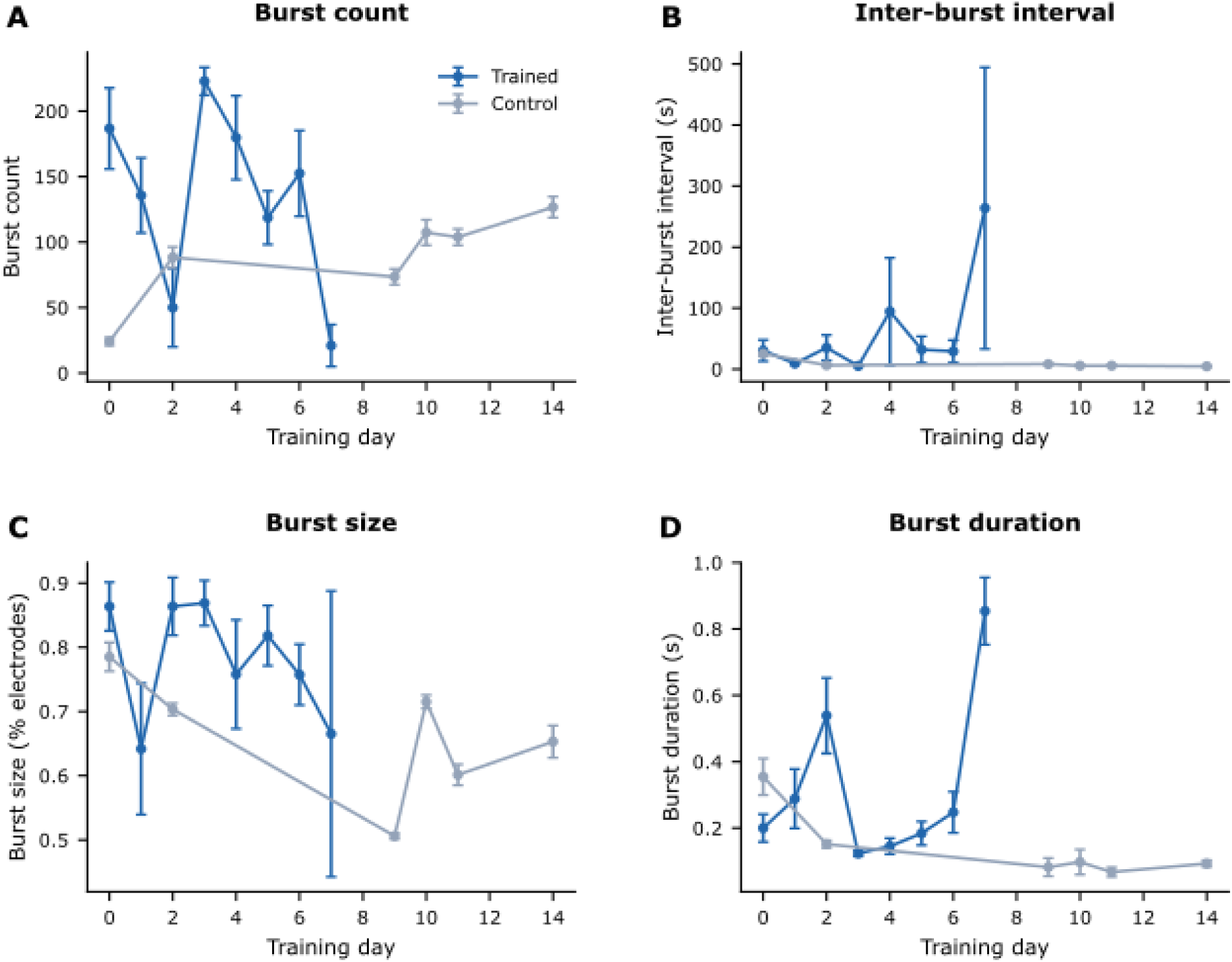
Burst properties over training days. Burst count (A), inter-burst interval (B), burst size (C), and burst duration (D) versus training day (pre-phase, mean and SEM), trained versus control. Descriptive companion to Figure 2.

**Figure S3.**
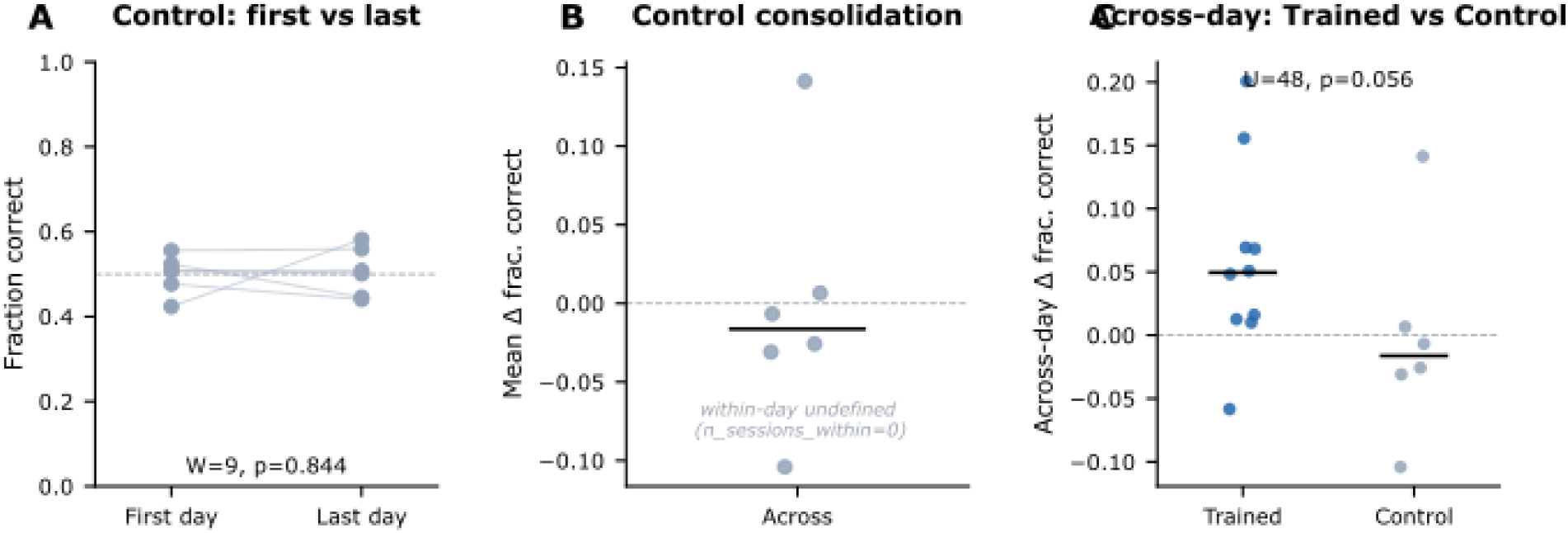
Control cultures provide a coin-flip null for the learning result. (A) Control cultures show no first-to-last improvement (paired Wilcoxon W=9, p=0.844). (B) Control across-day change; controls have no within-day data, so only the across-day term is defined. (C) Across-day change, trained versus control, the consolidation comparison missing from the main figure: the trained trend is higher but not significant (Mann-Whitney U=48, p=0.056; n=10 trained, 6 control).

**Figure S4.**
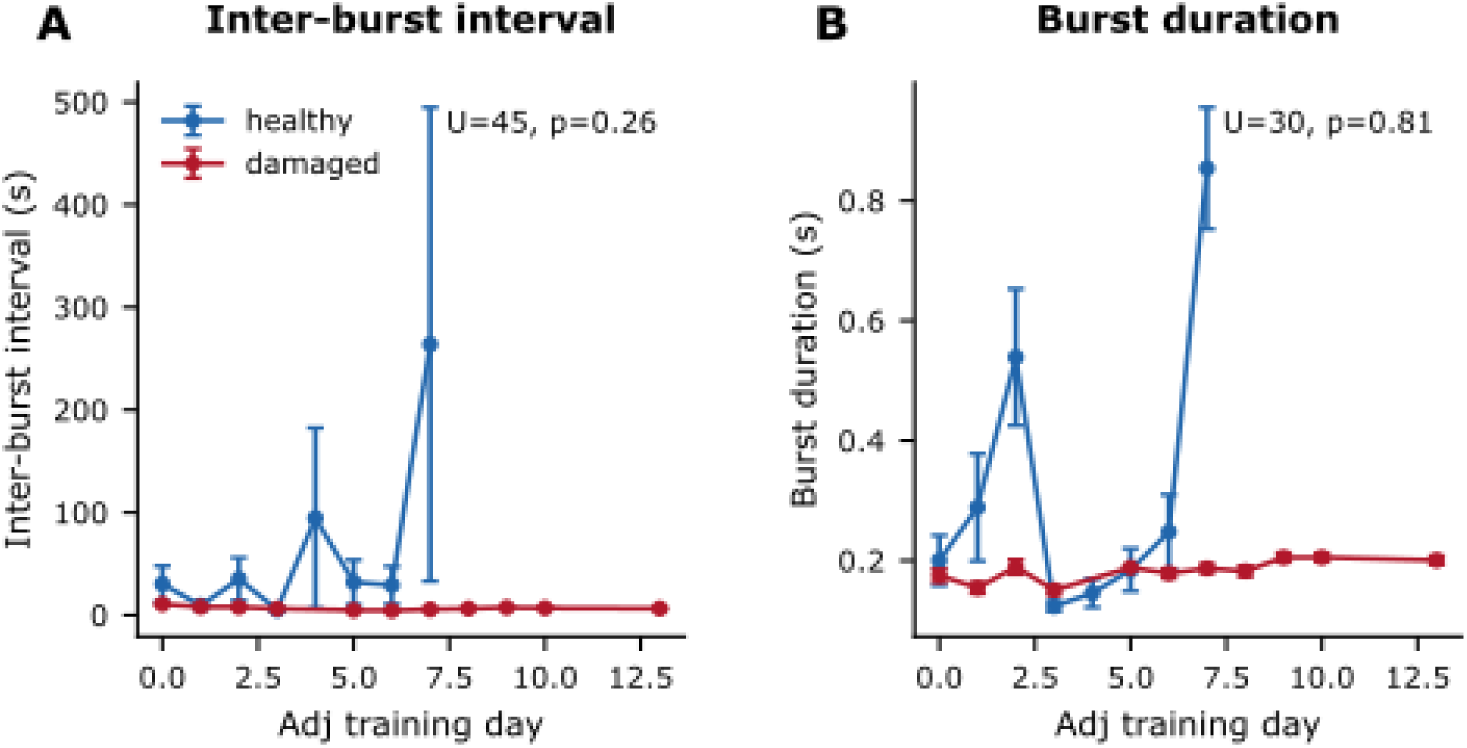
Damaged and healthy cultures share normal burst statistics. (A) Inter-burst interval and (B) burst duration over training days, healthy versus damaged; neither differs between groups (U=45, p=0.256 and U=30, p=0.808), supporting that damaged cultures burst normally.

**Figure S5.**
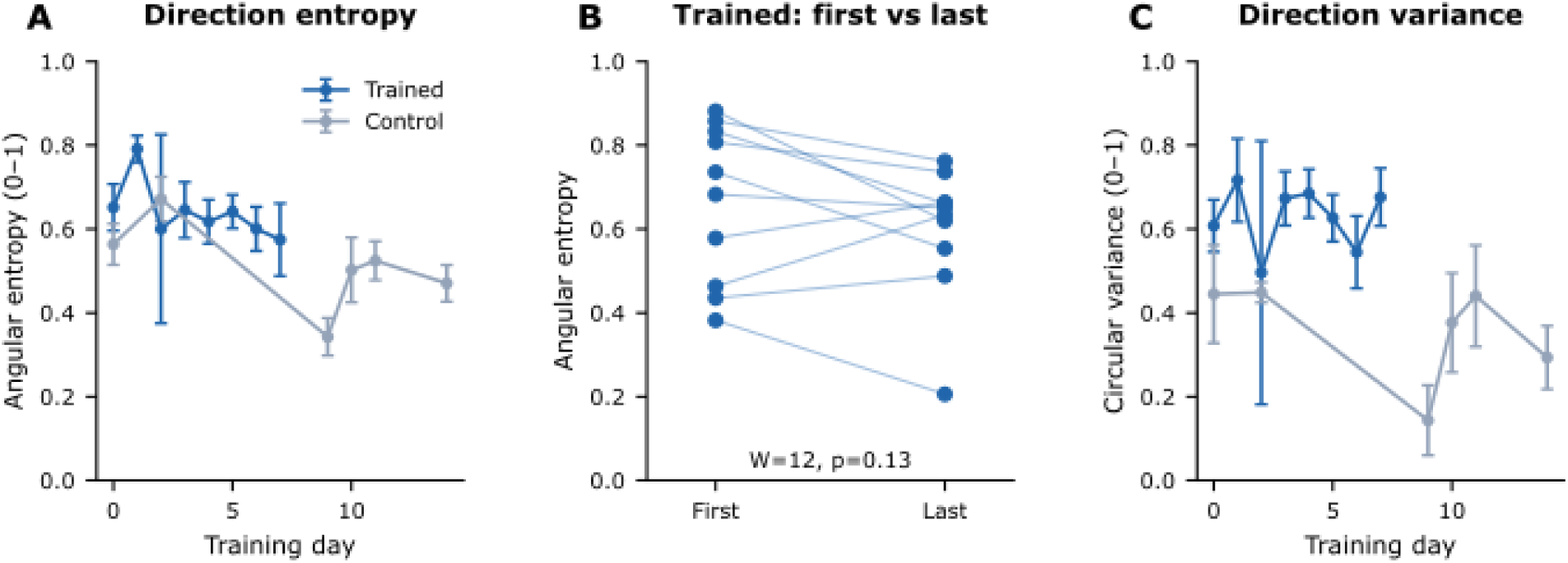
Training preserves the diversity of burst propagation directions. (A) Angular entropy of origin-to-peak vectors over training days, trained versus control. (B) Within-trained first versus last day of training for angular entropy, the null test that the direction repertoire does not change over training (angular entropy W=12, p=0.13, n=10; circular variance W=21, p=0.56). (C) Circular variance over training days, trained versus control. With the pooled per-culture comparison (angular entropy U=34, p=0.80; mean cosine U=41, p=0.80; circular variance p=0.80), training shifts the burst origin toward the goal without changing the diversity of propagation directions.

---

**Table T1.** Sensitivity of headline results to inclusion and stimulation filters. Headline tests recomputed under three filter settings (main; without the low-activity exclusion, retaining M07462; without the has_stim exclusion) to assess robustness. The stimulation-exposure measures reported in the main text (Figure 5D to F) are computed on unfiltered train-phase bursts and correspond to the third column. The values are tabulated below.

| Test | Main pipeline | Retaining<br>M07462 | No has_stim<br>exclusion |
| --- | --- | --- | --- |
| Fig 2B: learning, first vs last day (trained, Wilcoxon) | W=5, p=0.019 | W=5, p=0.019 | W=5, p=0.019 |
| Fig 2C: effect size, trained vs control (Mann-Whitney) | U=51, p=0.022 | U=51, p=0.022 | U=51, p=0.022 |
| Fig 4D: origin-shift vs peak-shift magnitude (trained, Wilcoxon) | W=3, p=0.010 | W=3, p=0.010 | W=3, p=0.010 |
| Fig 3C: recovery transition, trained vs control (Mann-Whitney) | U=46, p=0.026 | U=46, p=0.026 | U=46, p=0.026 |
| Fig 5F: stim fraction vs effect size (trained, Spearman) | rho=0.44,<br>p=0.200 | rho=0.44,<br>p=0.200 | rho=0.35,<br>p=0.328 |

**Table T2.** Baseline covariate sensitivity of the trained-versus-control effect size. Rank regression of the per-culture effect size on group, adjusting for one baseline burst property at a time (n=10 trained, 6 control). The group difference does not survive adjustment for baseline inter-burst interval or duration; because group and baseline differ together between groups recorded at different ages (Table 1), this indicates the contrast is confounded rather than that no learning occurred, consistent with treating the between-group comparison as supporting.

| Model | Group effect p |
| --- | --- |
| Unadjusted (group only) | p=0.017 |
| + baseline inter-burst interval | p=0.457 |
| + baseline burst duration | p=0.219 |

